# Decoding PDI diversity: insights into structure, domains, and functionality in sorghum

**DOI:** 10.1101/2025.06.06.658324

**Authors:** Carla F. López-Gómez, Marc T. Morris, Karen Massel, Millicent Smith, Peter Crisp, Gerhard Schenk, Ian D. Godwin

## Abstract

Proteins play indispensable roles in cellular function, acting as both structural components and catalysts for essential biological processes. Their proper folding into three-dimensional structures is critical for functionality. To ensure correct folding, proteins interact with chaperones and folding catalysts such as Protein Disulfide Isomerases (PDIs), which assist in the formation and rearrangement of disulfide bonds that stabilize proteins by linking cysteine residues. PDIs are part of the thioredoxin (TRX) superfamily and are characterized by a conserved CXXC motif that contributes to their redox potential. They exhibit isomerase and oxidoreductase activities, that enable them to rearrange and form new disulfide bonds. PDI family members in sorghum (SbPDI) present a broad and largely unexplored diversity in domain order, structure, and architecture between or even within species. To shed light on this diversity, we identified and characterized PDI family members in sorghum *in silico* to explore their domain architecture, three-dimensional structure and functionality.

**Author summary:** In this work, we explore how genomic and molecular tools can improve our understanding of the function and diversity of the protein disulfide isomerase (PDI) family in plants, using sorghum as a model and humans as a reference. By analysing domain architecture and predicted structures across the PDI family, we lay the groundwork for future studies investigating their roles in plant development and stress responses, including through targeted gene editing approaches. Although PDIs have been widely studied in humans, their structural and functional diversity in plants remains largely unexplored. With this study, I aim to help close this knowledge gap and highlight the structural differences of PDIs in plants.

## Introduction

Proteins are indispensable components in cells, not only because of their extensive structural roles but also since they catalyse the vast majority of essential biological functions. The reactions they catalyse maintain cellular metabolism and homeostasis, generate chemical messengers to regulate physiological processes, and play key roles in the immune system [1]. For proteins to successfully perform their specific functions with optimal efficiency, correct folding of their three-dimensional structure is crucial. In eukaryotic cells, most proteins are synthetised by ribosomes and translocated into the lumen of the endoplasmic reticulum (ER), where they encounter a series of chaperones, such as Protein Disulfide Isomerases (PDIs), which assist in their correct folding [2–4]. Disulfide bonds between two cysteine (Cys) residues establish a covalent linkage between different parts of a protein and contribute to both protein folding and stability. However, disulfide bonds formed in particular in the early stages of protein folding are often incorrect due to mis-paired cysteines residues [4]. PDIs facilitate the rearrangement of such incorrect disulfide pairings.

PDIs belong to the thioredoxin (TRX) superfamily, along with the peroxiredoxins (PRXs) and glutaredoxins (GRXs) [5, 6]. This superfamily is ubiquitously distributed in all living organisms and is characterised by a conserved CXXC catalytic motif (where C stands for cysteine and X for any amino acid side chain) [5, 7]. While bacterial representatives are generally small monomeric proteins, PDIs in most eukaryotes are multi-domain proteins resulting from the duplication or fusion of ancient monomeric TRX proteins [8]. Eukaryotic canonical PDIs generally consist of an N-terminal region (often containing a signal peptide of 15-30 amino acids in length), two TRX domains (both of which contain the CXXC motif, often referred to as motifs **a** and **a’**), two TRX-like domains (such as **b** and **b’**) that lack the motif, a linker (labelled **x**) and a disordered C-terminal region (Figure 1). The overarching PDI architecture can thus be described as **N**-**a-b-b’-x-a’-C**. The first complete crystal structure of a PDI (from yeast; PDB ID: 2B5E) was obtained at a resolution of 2.4 Å and illustrates the characteristic twisted “U” conformation that brings the two TRX domains in proximity [9]. Subsequent structures include human PDI (hPDI) in both the reduced and oxidized states at 2.9 Å and 2.5 Å resolution, respectively (Figure 1) [10]. Importantly, the two hPDI structures illustrate significant structural flexibility with a more closed reduced (PDB ID: 4EKZ) and more open oxidized (PDB ID: 4EL1) conformation [10].

**Figure 1.**
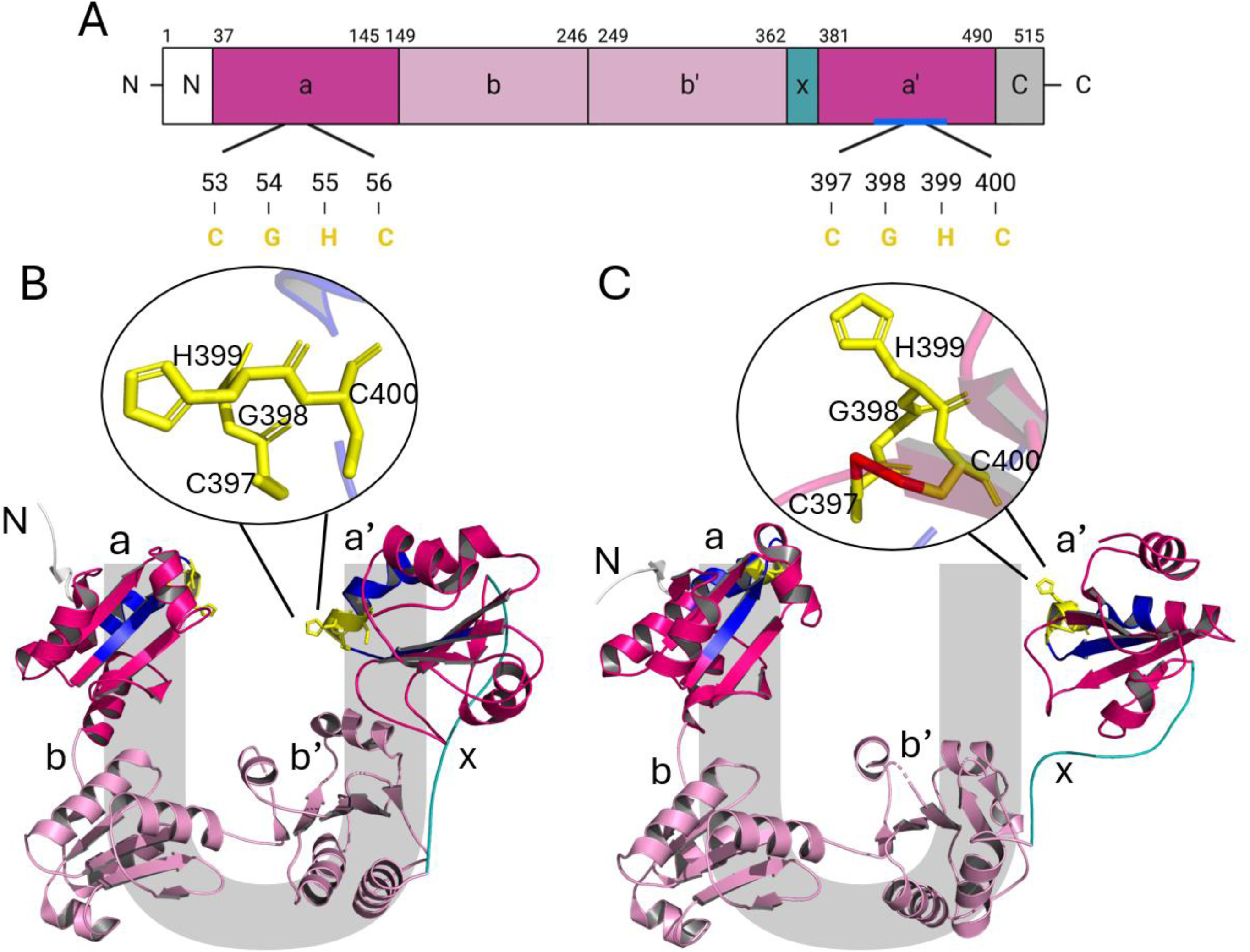
Domain teristic CXXC motif (CGHC in hPDI; yellow). PDI adopts two distinct conformations, depending on redox states of these motifs. In their reduced, non-bridged state they promote a closed conformation (B), in their oxidised, disarchitecture (A) and three-dimensional structure of a canonical PDI from human (B, C). From N-terminus to C-terminus: signal peptide (white), **a** and **a’** domain (magenta), **b** and **b’** domain (light pink), Linker **x** (cyan), and disordered C-terminal region (grey). The active sites are located in the TRX domains and contain the characulfide-bridged state an open one (C). The inset highlights the two active site states, reduced and oxidised (S atoms shows in red), respectively.

The TRX domains (**a** and **a’**) are approximately 100 amino acids in length and consist of four α-helices and four β-strands (Figure 1) [11, 12]. Within this motif, the two cysteine residues, when reduced and deprotonated, perform sequential nucleophilic attacks on the sulphur atoms of a misfolded protein (*i.e.,* the substrate) [13, 14]. The C-terminal **a’** domain is more flexible than the N-terminal **a** domain, which enables the former to accommodate a broader range of substrates [15]. TRX-like domains such as **b** and **b’** lost the CXXC motif and consequently their catalytic function after duplication of the ancestral TRX gene [11]. The N-terminal **b** domain is speculated to play a structural role while the C-terminal **b’** domain likely acts as the initial locus for substrate-binding in PDI as it is the only domain required for the binding of small peptides [16–18]. There is a hydrophobic substrate-binding pocket in the **b’** domain that likely interact with the substrate [17, 19].

The short regions located between each TRX and TRX-like domains act as linkers to provide molecular flexibility and enable conformational changes (between the open and closed states; Figure 1). These linkers generally consist of only a few amino acids, but Linker **x** between domains **b’** and **a’** is longer (up to 19 amino acids), enabling greater flexibility for the C-terminal end of PDI. Linker **x** is rich in proline and acidic amino acid residues [20, 21], and has been shown to reversibly cap and uncap the substrate-binding pocket of the **b’** domain through a tryptophan (Trp) residue in human PDIs [17, 22]. Notably, when PDI is oxidized without a bound substrate, the hydrophobic pocket remains capped, maintaining the CXXC motifs in the **a** and **a’** domains in their oxidized (open) states. However, in the presence of a bound substrate, the pocket becomes uncapped, leading to the reduction and closure of the active site (Figure 2) [18]. Finally, the disordered C-terminal region of the protein contains a KDEL signal peptide sequence which guides the polypeptide into the ER and a section rich in acidic residues. This region is likely to bind Ca^2+^ and hence may play a role in a signalling pathway [6, 23]. The C-terminal region may also mediate protein-protein interactions with other chaperones [6, 24].

**Figure 2.**
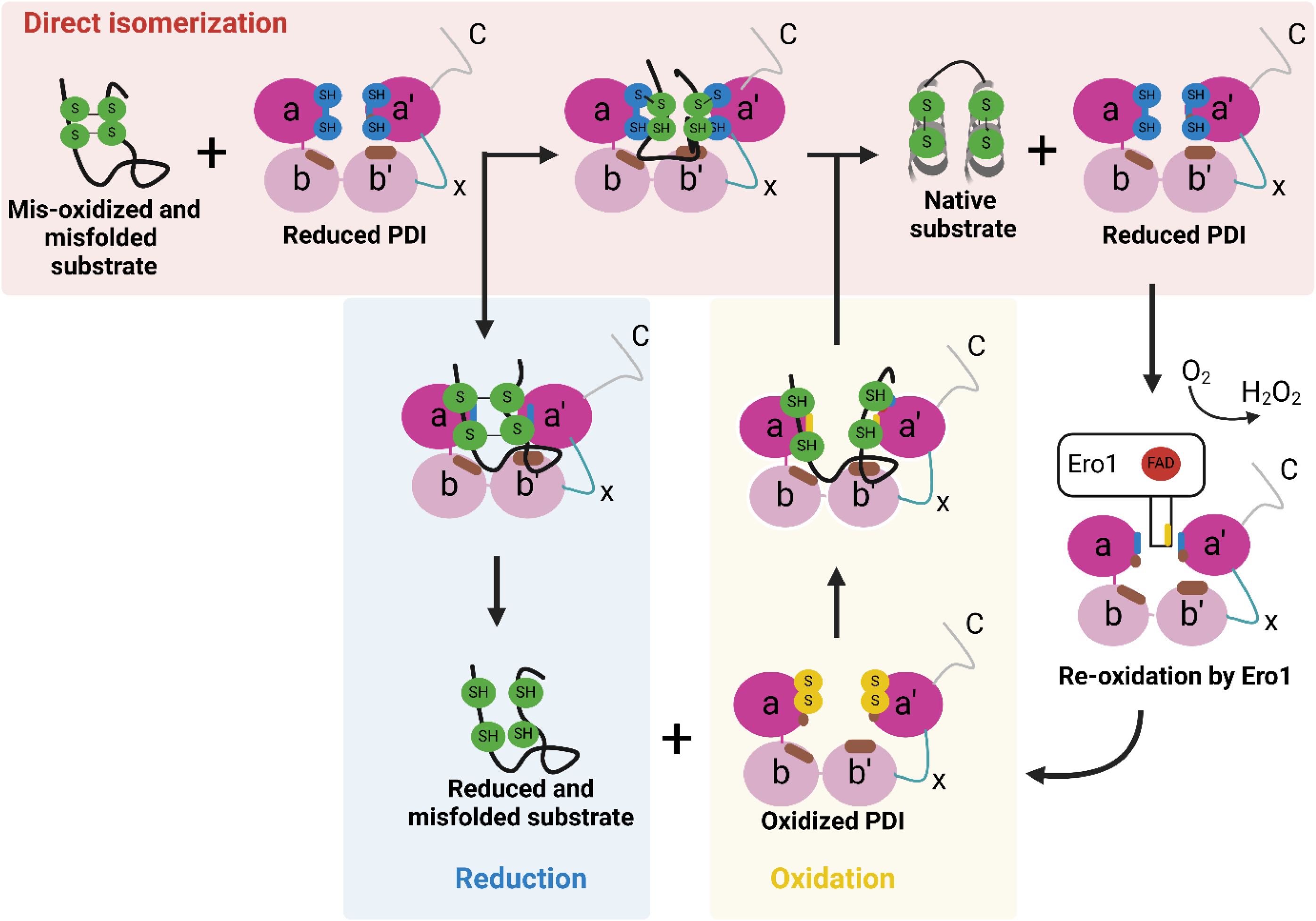
Scheme of the PDI-catalyzed reactions. PDI can perform both a direct isomerisation and an oxidoreductase reaction (see below for details).

Two distinct but synergistic catalytic mechanisms contribute to the chaperone function of PDIs, (*i.e.,* direct isomerisation and oxidoreductase activity) (Figure 2) [3, 4, 25]. In the direct isomerization, the misfolded protein (*i.e.,* the substrate) is bound to the reduced state of PDI and in a series of disulfide bond breakages and formations the properly folded (native) protein is formed while the active site is regenerated to its reduced state. In the oxidoreductase mechanism this concerted reaction is separated into two distinct steps. In the first step (reduction) the misfolded protein interacts with the reduced PDI, the mispaired disulfide bridges are reduced and broken, and the still misfolded protein is released. In the second step (oxidation) this protein binds to oxidised PDI and correctly paired disulfide bridges are formed while the reduced form of PDI is regenerated [14, 26]. The concentration of oxidised PDI in the lumen of the ER may be regulated *via* enzymes such as the flavin adenine dinucleotide (FAD)-dependent ER oxidoreductin (Ero1), which couple disulfide formation to the consumption of molecular oxygen [25, 27].

While canonical PDIs from human and microbial organisms have received most of the attention to date, the PDI family exhibits significant diversity in domain order and structure between or even within species. Plant PDIs, despite possessing both canonical and non-canonical representatives, have been poorly investigated to date. To address this gap, this study probes the architectural diversity of the PDI family in a major food and energy crop, sorghum.

## Results and discussion

### Domain architecture and predicted three-dimensional structures of SbPDIs

Members of the plant PDI family are classified into seven distinct subfamilies (I–VII), as defined by Houston et al. (2005). This study adopts those classifications to analyze the phylogenetic relationships between PDIs in plants. Specifically, within sorghum PDIs (SbPDIs), members of groups I, II and III exhibit the canonical domain architecture **a-b-b’-x-a’** (Table 1). In SbPDIs belonging to Groups IV and V, both the **a** and **a’** domains are located at the N-terminus followed by TRX-like domains (*i.e.,* the ERp29 and P5 domains, respectively; see Section 3.3). Notably, in Group IV, SbPDI2-1b has been identified in this study as part of the SbPDI family based on homology. However, it was not classified as such in the phylogenetic analysis by [7]. In SbPDIs that belong to Group VII, a single **a** domain is followed by the TRX-like domains **b** and **b’** to establish an **a-b-b’** arrangement. It is worth noting that no representative member from Group VI has been identified in sorghum. Figure 3 illustrates the domain architecture of the nine SbPDIs. Overall, the SbPDI family can be divided into two groups, those that exhibit the characteristic canonical architecture and those that differ (Table 1).

**Figure 3.**
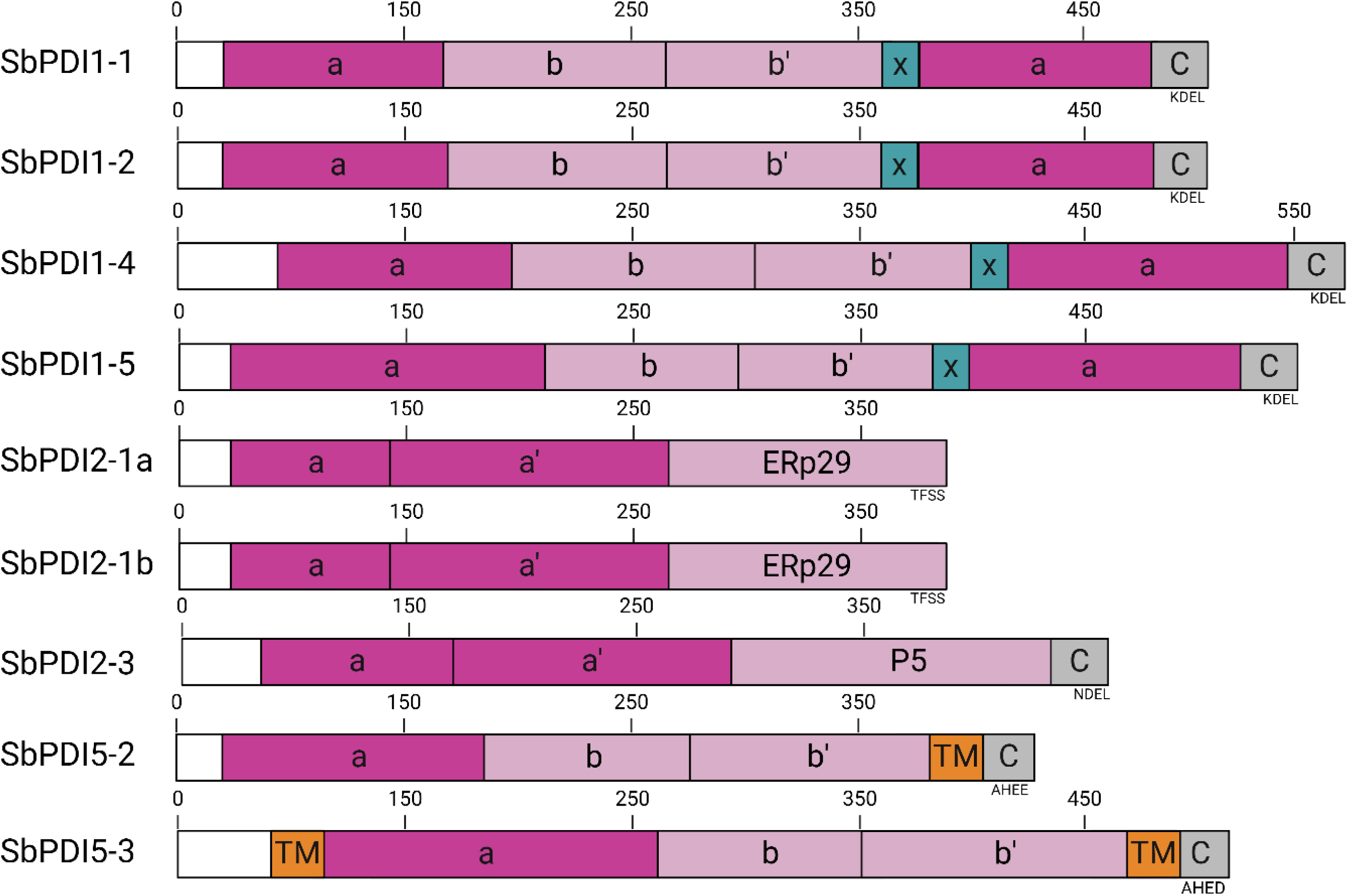
Domain architecture of SbPDIs. Note that TM stands for transmembrane regions, and **ERp29** and **P5** are TRX-like domains that share <15% sequence similarity to the more common **b** and **b’** domains and may have a different function (see Section 3.3) [42, 43].

**Table 1.** Classification of canonical and non-canonical PDIs identified in the sorghum genome. The phylogenetic groups, protein names, protein IDs (UniProt), protein length (in amino acids), and domain architecture are listed. Gene IDs (Phytozome) in order of appearance are Sobic.005G074400, Sobic.006G085700, Sobic.004G000400, Sobic.010G051100, Sobic.009G051600, Sobic.003G156400, Sobic.002G222200, Sobic.006G083400, Sobic.004G173600.

| Group | Protein name | Protein ID | Length | Domain architecture |
| --- | --- | --- | --- | --- |
| Canonical |  |  |  |  |
| I | SbPDI1-1 | A0A1B6PQR2 | 515 | **a-b-b'-x-a'** |
| I | SbPDI1-2 | A0A1B6PL31 | 518 | **a-b-b'-x-a'** |
| II | SbPDI1-4 | C5XRG2 | 572 | **a-b-b'-x-a'** |
| III | SbPDI1-5 | C5Z4V3 | 545 | **a-b-b'-x-a'** |
| Non-canonical |  |  |  |  |
| IV | SbPDI2-1a | C5Z0K4 | 367 | **a-a'-ERp29** |
| IV | SbPDI2-1b | A0A921RCE2 | 368 | **a-a'-ERp29** |
| V | SbPDI2-3 | C5X277 | 440 | **a-a'-P5** |
| VII | SbPDI5-2 | C5Y8P0 | 418 | **a-b-b'** |
| VII | SbPDI5-3 | A0A1Z5RN17 | 490 | **a-b-b'** |

The three-dimensional structures of the nine identified SbPDIs were predicted using AlphaFold 3 (Figure S1). The structural models of the four canonical SbPDIs could be validated by a comparison with the available structures of hPDI (Figure 1). However, there are currently no close reference structures to validate the accuracy of the structural predictions for SbPDI with a non-canonical architecture.

### Predicted structures of SbPDIs with a canonical architecture

SbPDI1-1, SbPDI1-2, SbPDI1-4, and SbPDI1-5 all share the canonical domain architecture **a-b-b’-x-a’** as exemplified by the three-dimensional structure of hPDI [10, 44]. Figure 4 shows the sequence identity at the amino acid level between SbPDIs and hPDI. While the highest degree of identity among these canonical SbPDIs is observed between SbPDI1-1 and SbPDI1-2 from the phylogenetic Group I (56 %), there is still significant identity between canonical SbPDIs from different groups (25-31 % identity). As expected, all canonical SbPDIs share higher identity among each other than with any of the non-canonical SbPDIs. Two isoforms of Group I PDIs are also present in rice, and *in silico* studies have led to the suggestion that they emerged from a gene duplication of a common ancestral gene [45]. Indeed, a widely accepted hypothesis for the evolution of PDIs involves a series of duplication and/or deletion events of an ancestral PDI protein containing a single TRX domain [8]. An alternative hypothesis suggests that the diversity in the PDI family may have been initiated by the fusion of two ancestral PDI proteins with single TRX domains [8]. To date, however, the evolutionary history of PDIs in general and SbPDIs in particular, remains obscure.

**Figure 4.**
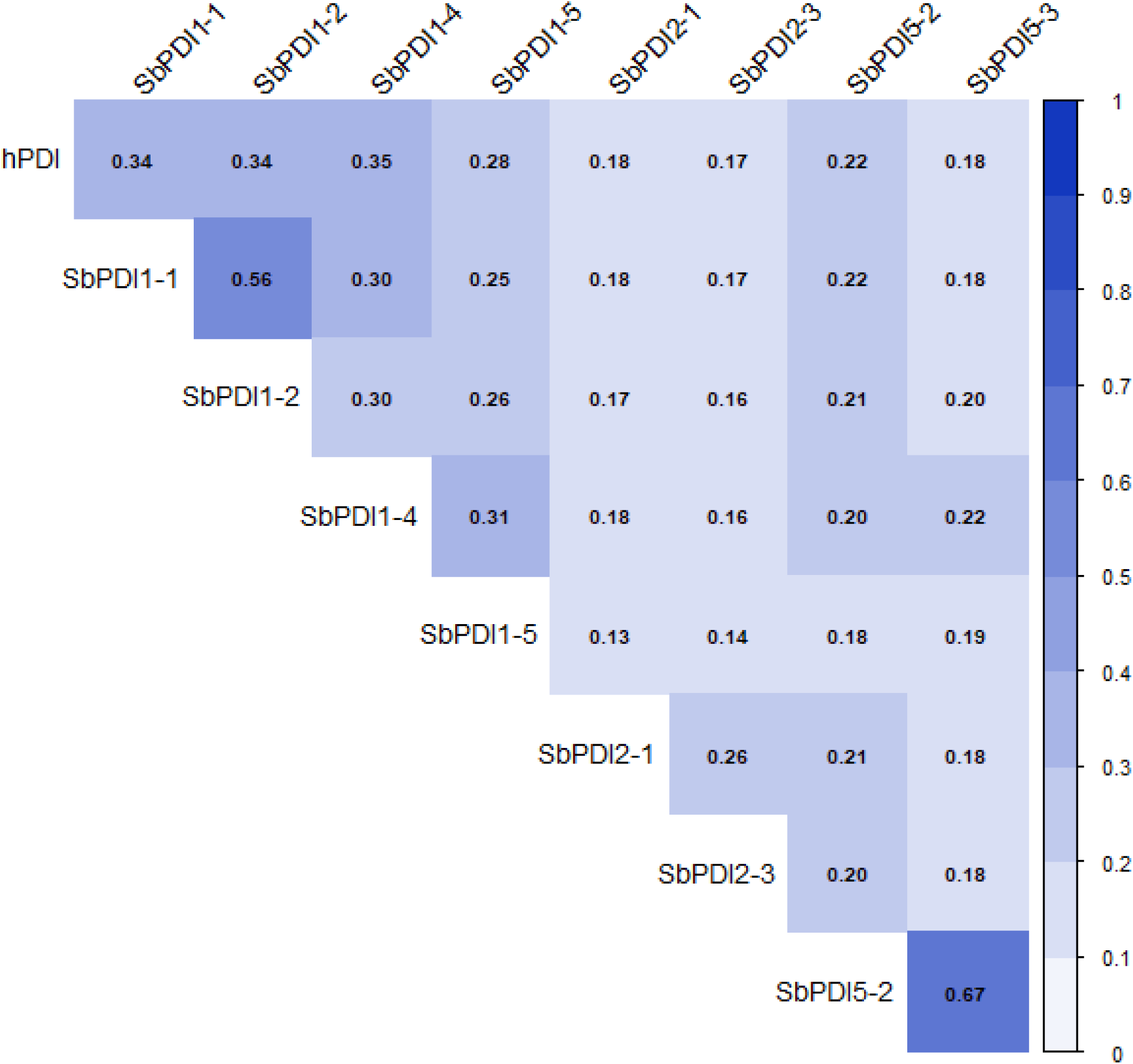
Heat map of the pairwise identity at the protein level between each SbPDI and hPDI. Note that SbPDI2-1 refers to SbPDI2-1a and SbPDI2-1b as both present the same identity scores.

Besides sharing very high sequence identity, the models of the four canonical SbPDIs appear structurally similar to hPDI, all showing the characteristic twisted “U” conformation (Figures 1 & A1). The models were superimposed onto the structures of both reduced and oxidised hPDI to estimate the root mean square deviation (RMSD; in Å) of structural pairs (Table 2). For SbPDI1-1, the structural alignment is better with the reduced, closed form of hPDI (4.4 Å), while for the other three canonical SbPDIs a better alignment was achieved with the oxidised, open form of hPDI (4.0 Å – 5.7 Å). In particular, SbPDI1-4 shares closer structural similarity to the oxidised human enzyme (4.0 Å). Noteworthy, in SbPDI1-4 the catalytic centre is modelled in its reduced (-SH) state. The differences between the phylogenetically closely related SbPDI1-1 and SbPDI1-4 [7] are interesting as they may illustrate two catalytically distinct states that invoke significant structural flexibility throughout the reaction mechanism (Figure 2). While functional insight gained from such predicted structural models need be viewed with caution, it is nonetheless possible that canonical plant PDIs may employ a mechanism that varies from that of their better studied mammalian counterparts.

**Table 2.** Pairwise superimposition between SbPDIs and either reduced or oxidised hPDI. For the comparison the structures were superimposed using PyMOL software, including only the backbone atoms from the pairwise comparisons to calculate the RMSD values.

|  | Reduced hPDI |  | Oxidized hPDI |  |
| --- | --- | --- | --- | --- |
|  | RMSD (Å) | Backbone atoms | RMSD (Å) | Backbone atoms |
| Canonical |  |  |  |  |
| **SbPDI1-1** | 4.4 | 2634 | 6.7 | 2747 |
| **SbPDI1-2** | 6.0 | 2701 | 5.7 | 2615 |
| **SbPDI1-4** | 6.4 | 2876 | 4.0 | 2264 |
| **SbPDI1-5** | 8.7 | 2666 | 5.4 | 2325 |
| Non-canonical |  |  |  |  |
| **SbPDI2-1a** | 18 | 1555 | 21.0 | 1560 |
| **SbPDI2-1b** | 17 | 1536 | 20.1 | 1542 |
| **SbPDI2-3** | 16.8 | 1491 | 16.9 | 1497 |
| **SbPDI5-2** | 6.9 | 1820 | 8.2 | 1880 |
| **SbPDI5-3** | 6.7 | 1862 | 8.0 | 1844 |

SbPDI1-5 is part of the phylogenetic Group III, and in contrast to the other known canonical SbPDIs, it displays a variation in its active site motif. Instead of the commonly found CGHC motif, its two TRX domains use the CERS and CVDC motif as previously reported in rice and wheat [45], suggesting a variation of the redox potential of these active sites. As mentioned above, PDIs require both cysteine residues in the active site motif to perform their oxidase function, but the N-terminal Cys alone is sufficient for isomerase activity [13]. Therefore, the CERS site of SbPDI1-5 is likely to perform only isomerase reactions whereas the CVDC site is expected to be able to employ both mechanistic strategies (Figure 2). PDIs with divergent motifs are suspected to act as retention or redox agents in the ER instead of chaperones, in a similar way to ERp44 in soybean [46–48]. ERp44 (UniProt ID: Q9BS26) possesses a TRX domain with a non-canonical CXFS motif similar to the CERS motif of SbPDI1-5. In mammals, ERp44 captures improperly folded proteins through its exposed cysteine residue and returns those proteins into the ER. In fact, ERp44 is involved in retaining Ero1, which is in turn responsible for regenerating the reduced PDI (Figure 2) [25]. SbPDI1-5 and ERp44 share only 15 % pairwise sequence identity overall, whilst their non-canonical TRX domains share 27 % identity. At the structural level, ERp44 possesses one domain less (**a-b-b’**) than SbPDI1-5 (Table 1), and its structure thus differs from the canonical PDI conformation. However, SbPDI1-5 is the only member in sorghum with a CXXS motif and may perform a similar function to ERp44 despite significant differences in their architecture. SbPDI1-5 may thus be a potential isoform of ERp44, for which an equivalent protein has not been previously identified in plants.

### Predicted structures of SbPDIs with a non-canonical architecture

SbPDI2-1, SbPDI2-1a, SbPDI2-3, SbPDI5-2 and SbPDI5-3 show significant differences from the canonical architecture (Table 1; Figure 3) and share less than 22% sequence identity with canonical PDIs (Figure 5). In all non-canonical SbPDIs, the C-terminal KDEL signal for retention in the ER present in their canonical counterparts is replaced by sequences such as TFSS, NDEL, IHEE or AHED (Figure 3). A similar observation was previously reported for homologous PDIs from rice and wheat [45]. The function of the NDEL motif appears to be same as that of KDEL, but it is unknown, to date, whether the other motifs bestow a similar function.

**Figure 5.**
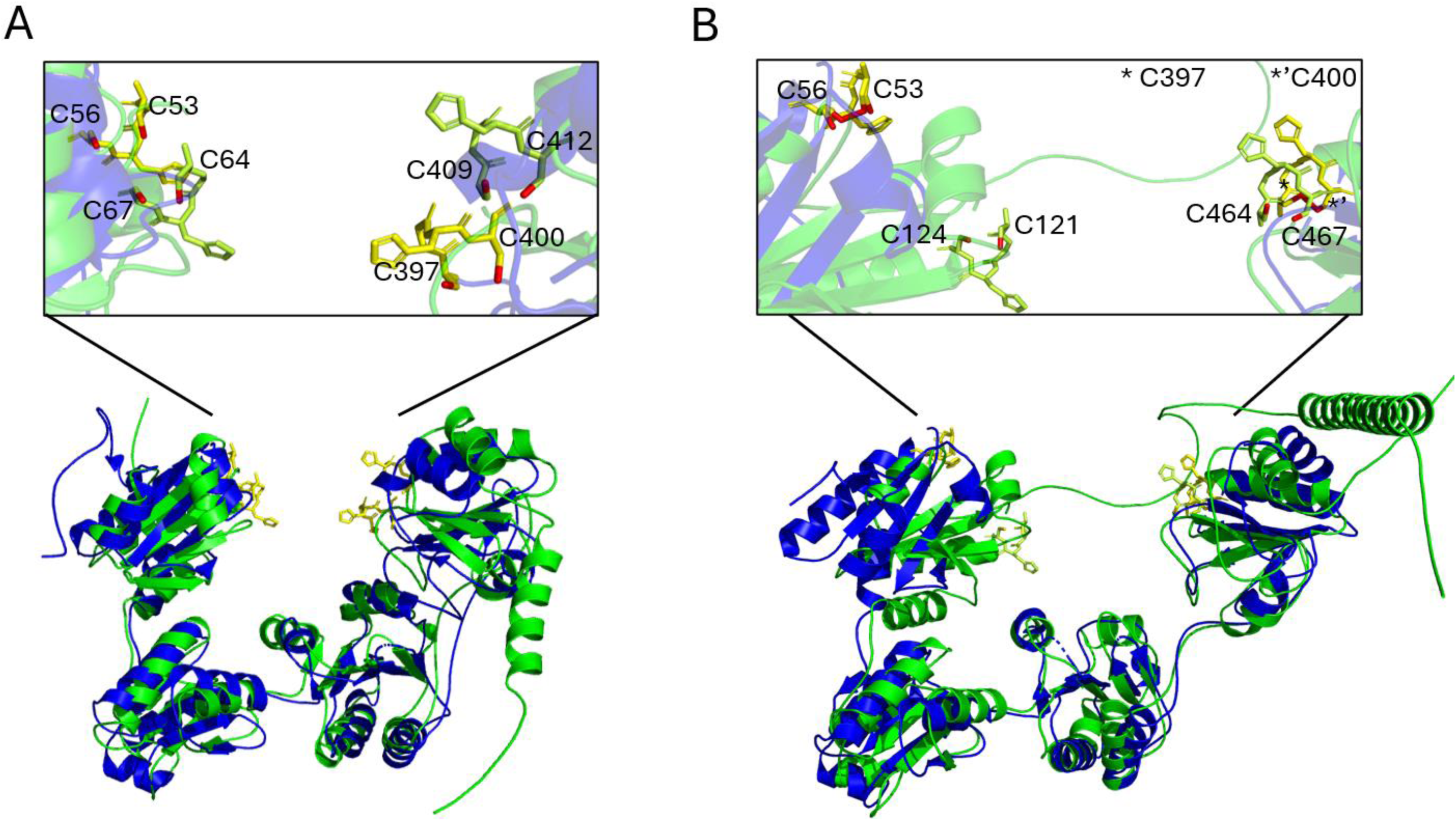
Superimposition of selected structural models of SbPDIs with hPDI. (A) SbPDI1-1 (green) superimposes with an RMSD of 4.4 Å with the reduced/closed form of hPDI (blue). (B) SbPDI1-4 (green) aligns better with the oxidized/open form of hPDI (blue) with an RMSD of 4.0 Å. For both superimpositions, the CGHC motifs (yellow for hPDI, lime for SbPDI) and S atoms (red) are shown. Note that N- and C-terminal regions have been truncated to simplify the display.

SbPDI2-1a and SbPDI2-1b belong to the phylogenetic Group IV and share 81 % of pairwise identity (Figure 4), the highest observed for SbPDI family. It is the first time that SbPDI2-1b is classified within the SbPDI family as a potential duplicate or homoallele of SbPDI2-1a. They both contain two TRX domains followed by a C-terminal ERp29 domain in an **a**-**a’**-**ERp29** arrangement as previously suggested for wheat and rice PDIs [45]. Their structural models align poorly with both reduced and oxidised hPDI (Table 2). Interestingly, the ERp29 protein (UniProt ID: P30040), from which the domain name is derived, is an ER-resident protein that also has high levels of expression in the extracellular medium in mammals [49]. In mammals, the monodomain ERp29 acts as a secretion factor and it is co-upregulated with ER chaperones and PDIs [50]. The two active TRX domains in SbPDI2-1a and SbPDI2-1b are expected to promote effective oxidoreductase and isomerase activities (see Introduction). Furthermore, the ERp29 domain, together with a 15-amino acid linker (G258-L273) that connects this domain to the **a’** domain, may enhance the functionality of SbPDI2-1a and SbPDI2-1b as chaperone and possibly also facilitate cellular trafficking. In addition, both SbPDI2-1a and SbPDI2-1b lack the KDEL signal for retention within the ER, allowing them to move freely in and out of this organelle. In the homologous protein from Arabidopsis (PDIL2-1), the C-terminal ERp29 domain has been shown to play an important role in embryo sac maturation and pollen tube guidance [51, 52]. Taken together, we propose that SbPDI2-1a and SbPDI2-1b may act as escort chaperone proteins, assisting in the transport, localization and proper folding of other proteins.

The Group V SbPDI2-3 has an architecture similar to that of SbPDI2-1a, but with a **P5** instead of an **ERp29** domain in an **a-a’-P5** arrangement (Table 1; Figure 3). The two non-canonical SbPDIs share 26% identity (Figure 5). The **P5** domain alone shares 43.2% identity with the C-terminal region of the human P5 protein (UniProt ID: Q15084), also known as ERp5, which has an **a-a’-P5** architecture [53, 54]. Interestingly, the C-terminal TRX-like domain of a soybean PDI also shares a similar sequence identity with the corresponding domain in ERp5 [42]. In ERp5, the **P5** domain enables the formation of complexes with several ER chaperones and folding factors to inhibit aggregation of misfolded proteins [55]. In addition, ERp5 has been reported to interact with other PDIs forming larger complexes in mammals [39]. ERp5 has a variation in the redox site, with three additional amino acids separating the essential Cys residues (*i.e.,* CX_5_C). This motif may still facilitate redox processes [56]. Interestingly, ERp5 forms a homodimer using a leucine-valine (Leu-Val) adhesive motif (V124, L128, L131, V135, L139) located in the **a** domain [53]. This motif is also present in the **a** domain of SbPDI2-3 (V125, L129, V132, L136, L140). Within the SbPDI family, the Leu-Val adhesive motif seems to be unique to SbPDI2-3. Other SbPDIs possess shorter Leu-Val-rich regions (1-3 residues) in the same position within the **a** domain, but *in silico* predictions suggest that only SbPDI2-3 may form dimers (Figure 6). Although the AlphaFold-derived pTM score for the SbPDI2-3 dimer model is relatively low (∼0.37), a similar score was obtained for the homodimer of ERp5, for which crystallographic evidence for dimerization is available [39]. Based on the models, we propose that SbPDI2-3 is likely homodimeric, presumably performing a function equivalent to that of ERp5.

**Figure 6.**
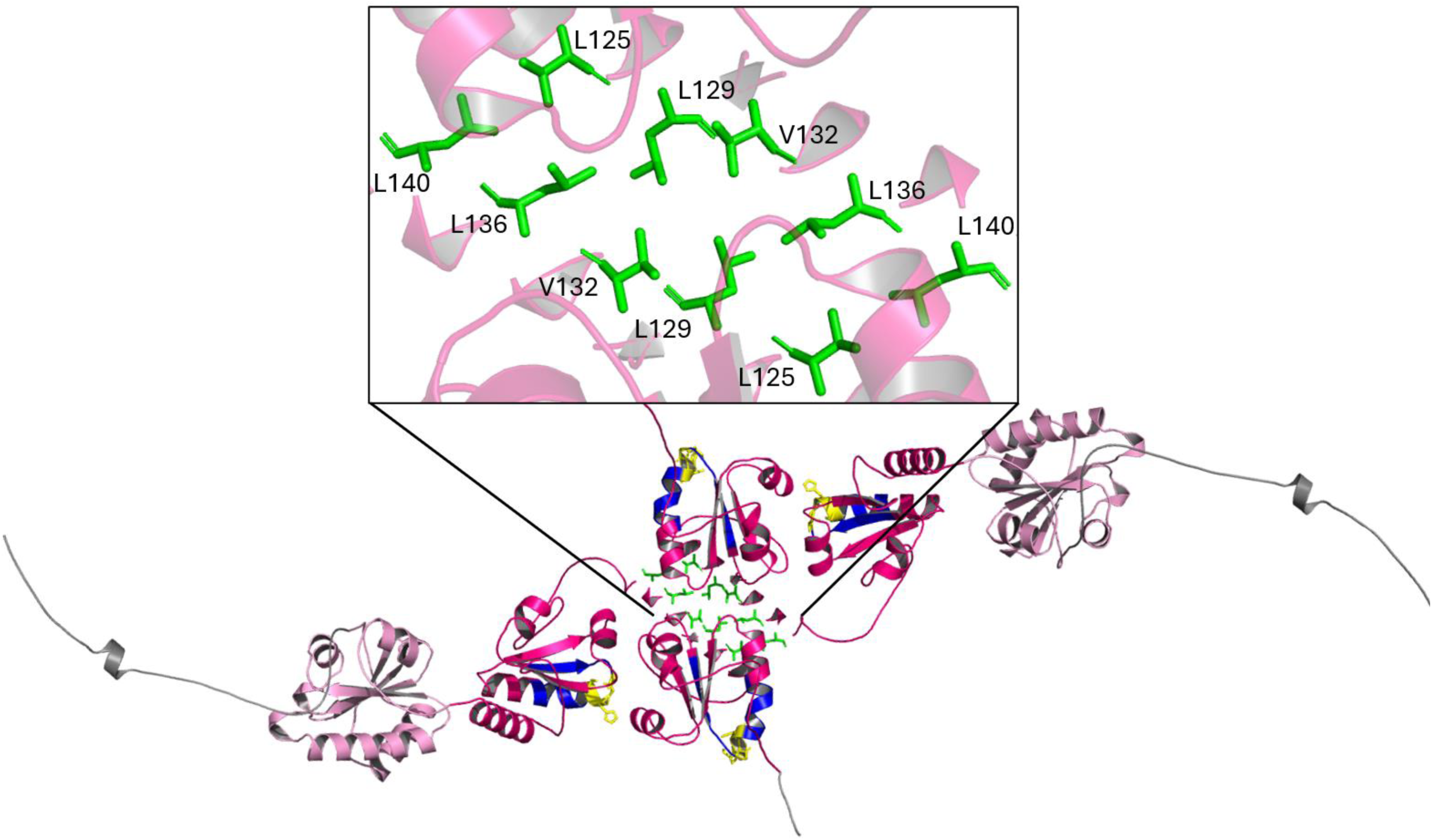
Predicted dimerization of SbPDI2-3. From the N-to the C-terminus: TRX domains **a** and **a’** (dark pink), active site (blue), redox site (yellow), **P5** domains (light pink), C-terminal region (grey), Leu/Val adhesive motif (green). Note that the N-terminal regions have been truncated to simplify the display.

Finally, SbPDI5-2 and SbPDI5-3 from Group VII, share significant identity with each other (66.7 %; Figure 4) and have an architecture that resembles the N-terminal half of canonical PDIs (*i.e.,* **a-b-b’**; Table 1; Figure 3). Consequently, it is not surprising that the structural models for these two proteins superimpose well on the structure of hPDI, in particular the reduced form (∼6.8 Å; Table 2). It has been previously suggested that their homologs in rice and wheat might be duplicates or homoalleles [45]. According to DeepTMHMM predictions, both SbPDI5-2 and SbPDI5-3 present transmembranal regions, thus suggesting that they are ER membrane-bound. Using these predictions, a structural model of SbPDI5-2 within the membrane has been created (Figure 7).

**Figure 7.**
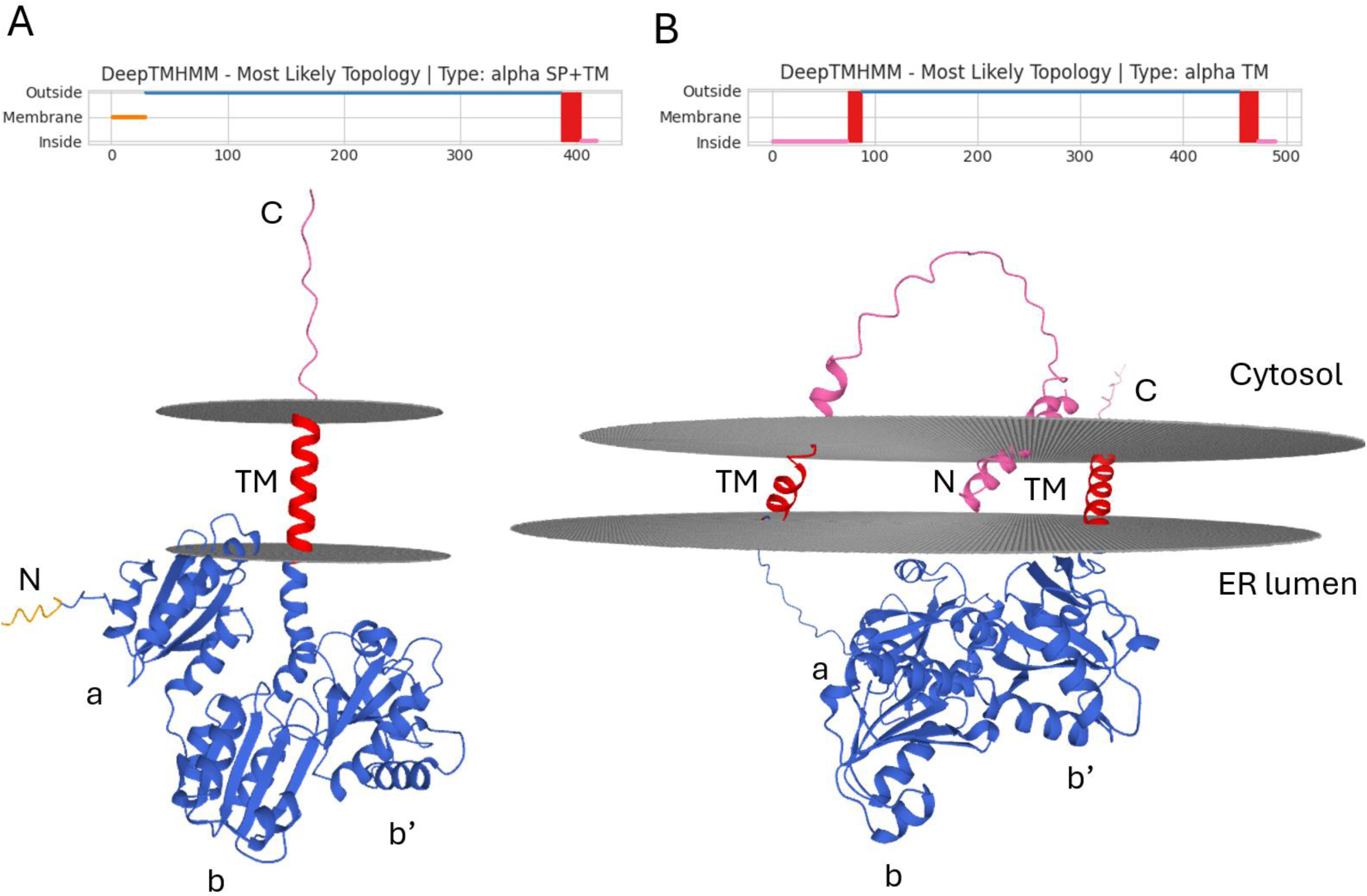
Predicted structure of transmembrane regions in (A) SbPDI5-2 and (B) SbPDI5-3 using DeepTMHMM software. Note that the α-helix inside the membrane is a model inaccuracy resulting from the challenge of predicting transmembrane proteins.

In SbPDI5-2, this transmembrane region may be located at the N-terminus, while SbPDI5-3 may have both N- and C-terminal transmembrane domains (Figures 8 & A1). It should, however, be noted that all structural predictions were made under conditions that mimic a cytosolic environment, and hence predictions about transmembrane conformations need to be viewed with caution. A thioredoxin-related transmembrane protein family (TMX) has previously been reported in mammals but not in plants [57]. Within this family, uniquely TMX3 (UniProt ID: Q3B7X1) presents the CGHC motif, one transmembrane domain, and a similar domain architecture (**a-b-b’**) to SbPDI5-2 and SbPDI5-3. The human TMX3 shares 20% protein sequence identity with both SbPDI5-2 and SbPDI5-3. Nevertheless, the structural superimposition of the AlphaFold models of TMX3 and SbPDI5-3 is better (3.8 Å for 1421 backbone atoms; Figure S2) than using SbPDI5-2 (5.1 Å for 1781 backbone atoms; Figure S2). TMX3 is involved in the folding of transmembrane proteins and acts as a cofactor for the functional expression of several receptors [57]. This suggests that SbPDI5-2 and SbPDI5-3 may constitute plant counterparts to the mammalian TMX protein family, which has not been identified or characterized in plants.

**Figure 8.**
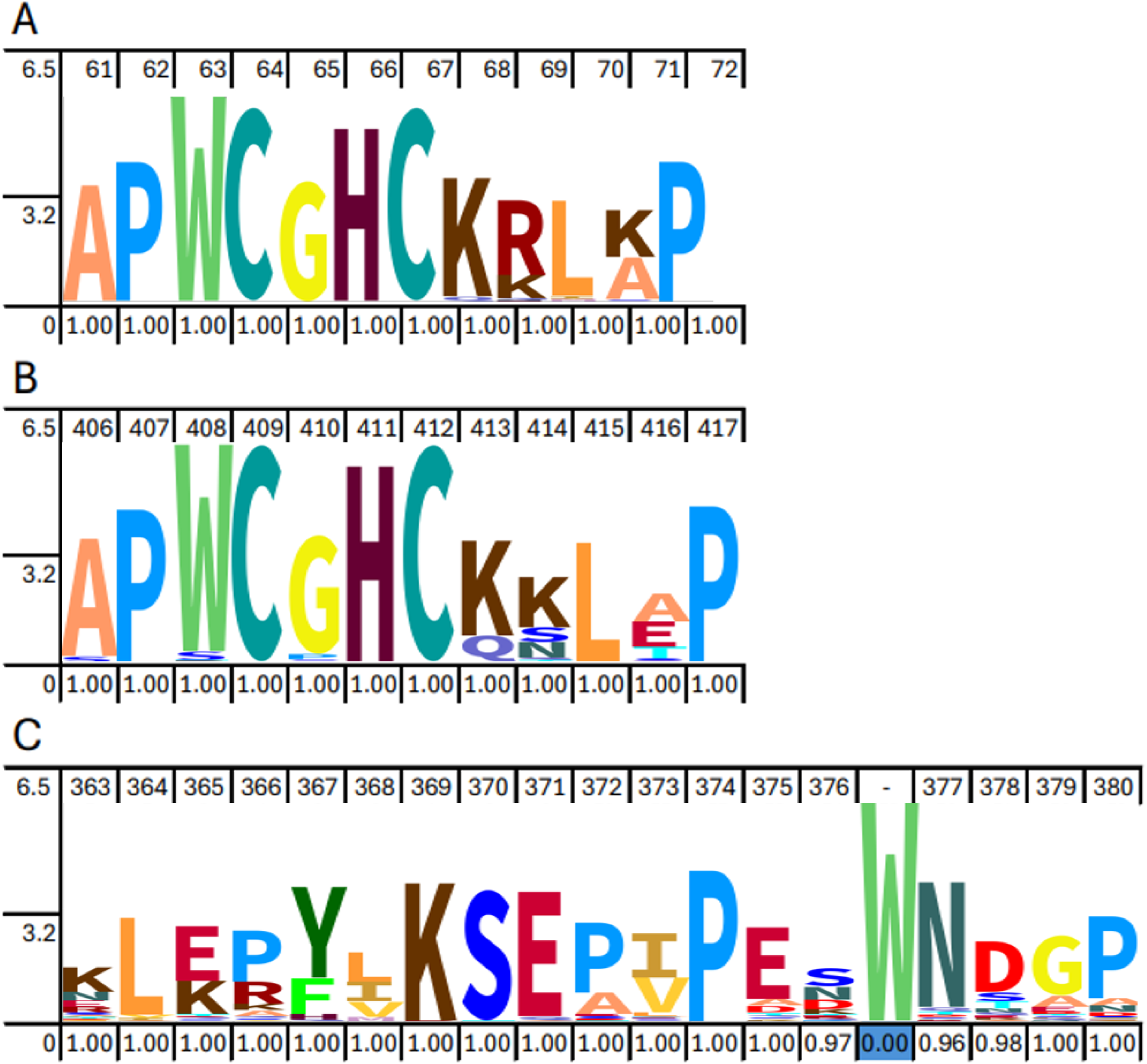
HMM logo of conserved active sites among 1,355 different PDI sequences from Eukarya. (A) Active site of the N-terminal (*i.e.,* **a** domain) and (B) C-terminal TRX domain (**a’** domain). (C) Linker **x** between domains **b’** and **a’** in canonical PDIs. Note that a gap (-) is present due to the absence of the tryptophan residue.

### Conserved key amino acids in the SbPDI family

Apart from the differences in domain architecture and their three-dimensional structures, protein families frequently share highly conserved and functionally essential amino acids, even across different species. In this section, conserved residues are highlighted for SbPDIs. For simplicity, amino acid positions have been numbered using the sequence of the canonical SbPDI1-1 as a template. Since the CXXC motifs in the **a** and **a’** domains are highly conserved in most SbPDIs, it is expected that they are all able to promote efficient oxidoreductase and isomerase activities (see Introduction). Other relevant conserved residues include Trp63 and Trp408 (Figures 9A & 9B), which precede the CXXC motif in the **a** and **a’** domains, respectively, and which are exposed to the solvent based on the structure of hPDI and the models of the SbPDI (Figure S3). In hPDI, this tryptophan has been linked to the thermostability of this enzyme [21], and thus it is likely that it performs a similar role in SbPDIs. Together with the conserved residues Ala61/Asp94 and Ala406/Asp438, Trp63/Trp408 form a structural cluster that was previously identified in hPDI, where it is proposed to stabilize its multi-domain architecture [12]. Indeed, the replacement of this Trp by an Ala in yeast PDI causes an interchange of the **a** and **b** domains changing the architecture to **b-a-b’-a’**, and resulting in an inactive, non-functional protein [58].

Two proline residues (Pro62/Pro72, Pro407/Pro417) immediately up- and downstream from the catalytic CXXC motifs in the **a**/**a’** domains are also highly conserved among different plant species, including SbPDIs, except after the CVDC motif in SbPDI1-5 (Figures 9A & 9B). Pro62/Pro407 introduce a conformational kink in the α-helix of the active site that separates the CGHC motif, located at the amino end of the same α-helix, from the following β-sheet [9]. In bacteria, the mutation of Pro62/Pro407 to Ala destabilizes the TRX structure, bringing the active site cysteines closer together, which in turn reduces the reactivity of each active site [9].

Other conserved residues in the TRX domains form hydrophilic and hydrophobic patches located proximal to the active site and are likely involved in protein-protein interactions as shown in hPDI-Ero1 docking studies. These include a phenylalanine-rich hydrophobic patch in the substrate-binding pocket of the **b’** domain, which is also present in SbPDI1-1 (F278, F300, F322) [27, 59]. This hydrophobic patch is exposed to the solvent in hPDI, however, only F322 is highly conserved across eukaryotes, including SbPDIs (File B1).

As mentioned above, Linker **x** (Figure 8C), at least in the canonical hPDI, may play an important role in the catalytic mechanism by reversibly capping the substrate-binding pocket, which also governs the redox state of the catalytic site. A tryptophan residue has been reported to be crucial in this process [17]. In hPDI, Linker **x** (located between the **b’** and **a** domains) contains 19 amino acids and, but only 18 in SbPDI1-1, SbPDI1-2 and SbPDI1-4, and 17 amino acids in SbPDI1-5. In non-canonical SbPDIs, Linker **x** is not present (Figure 3). Importantly, the tryptophan residue is not conserved in PDI homologs across different eukaryotic species. Instead, the structurally equivalent position appears to be occupied by a proline (Pro374) in canonical SbPDIs. This proline residue is conserved in eukaryotic species (File B1, Figure 8C). It is currently unknown whether Pro374 plays a similar role as Trp in hPDI in the reversible capping of the substrate-binding pocket.

## Methods

### Comparative structural analysis

PDI protein sequences from sorghum (SbPDI) were extracted from the Phytozome database [28] from *Sorghum bicolor* v3.1.1 genome and named following the rice PDI nomenclature. SbPDI family members include SbPDI1-1 (Sobic.005G074400), SbPDI1-2 (Sobic.006G085700), SbPDI1-4 (Sobic.004G000400), SbPDI1-5 (Sobic.010G051100), SbPDI2-1a (Sobic.009G051600), SbPDI2-1b (Sobic.003G156400), SbPDI2-3 (Sobic.002G222200), SbPDI5-2 (Sobic.006G083400) and SbPDI5-3 (Sobic.004G173600). Protein domains and active site residues were identified using the InterPro database [29], which integrates different databases to classify proteins into families and to predict domains and important sites within their protein sequences.

Models of the three-dimensional structure of each SbPDI member were predicted using AlphaFold 3 [30] and analysed with PyMOL 3.1 [31] based on the electrostatic surface potential and hydrophobicity index. A set of Python scripts was developed to assist with the analyses conducted in PyMOL 3.1. The accuracy of the predicted three-dimensional structures of canonical SbPDIs was validated by superimposing these models onto the experimentally determined crystal structures of hPDIs (Figure 1) using PyMOL 3.1. The same process was used to analyse the potential protein homologs of SbPDIs identified in this study, including ERp44 (UniProt ID: Q9BS26), ERp29 (UniProt ID: P30040), P5 or ERp5 protein (UniProt ID: Q15084), and TMX3 (UniProt ID: Q3B7X1).

The superimpositions for each SbPDI model and hPDI were made using PyMOL 3.1 software. The RMSD values for superimposed backbone atoms were calculated using the same software.

### Analysis of the active site conservation and pairwise identity

To study the conservation of the active site, FASTA sequences of the homologous gene family of SbPDI1-1, SbPD2-1a, SbPDI2-1b and SbPDI5-1 from 1,355 different eukaryotic species were extracted from the Plaza 5.0 [32] database and compared in a multiple sequence alignment (MSA) using MUSCLE software [33] with default parameters in Geneious Prime 2.0 [34]. The MSA file was curated, removing those sequences without at least one CXXC conserved domain, and used to create a consensus sequence logo of the active site using the Skylign database [35]. The raw file (File B1) containing all FASTA sequences can be accessed in [36]. Using the same alignment protocol as above, protein sequences for each SbPDIs and hPDI were extracted and aligned in a pairwise manner. The obtained protein sequence identity scores between each SbPDI and hPDI (0-1) were used to create a heatmap using RStudio [37].

### Prediction of the oligomeric state of SbPDI2-3

The protein sequence of SbPDI2-3 (Sobic.002G222200) was extracted from the Phytozome database [28] and the sequence of P5, a human PDI that occurs in dimeric form, was extracted from UniProt database [38]. For SbPDI2-3, based on the presence of the Leu-Val adhesive motif contact, a dimer model was generated using AlphaFold 3 [30] and analysed using PyMOL 3.1 [31]. The predicted dimer was validated by regenerating the P5 dimeric structure with the same approach and comparing this model with the experimental structure of that protein [39]. The obtained pTM scores were ∼0.37 for all the predictions, including SbPDI2-3 and P5 dimerizations.

### Prediction of the transmembrane regions of SbPDI5-2 and SbPDI5-3

The protein sequences of SbPDI5-2 and SbPDI5-3 were used to predict transmembrane regions using DeepTMHMM [40] software. Structural models within the membrane were then generated using MembraneFold [41].

## Conclusion

This study has highlighted the diversity of the PDI family in sorghum. Specifically, both canonical and non-canonical PDIs are abundant in sorghum (Table 1; Figure 3). The canonical variants are structurally closely related to their counterpart in human (Table 2; Figure S1), with most catalytically relevant residues well conserved (Figure 8). Nonetheless, plant PDIs, and possibly other eukaryotics, may employ a mechanism distinct from that used by the human counterpart. This distinction is suggested by differences in some key residues (*i.e,* the absence of a catalytically relevant Trp in hPDI, whose function is likely assumed by a Pro in canonical SbPDIs; Figure 8C) and possibly the impact of substrate-binding on the redox state of the catalytic CXXC motif (Figure 5). Furthermore, the presence of several non-canonical PDIs in sorghum also indicates that the portfolio of possible functions for these proteins goes beyond governing proper folding of proteins. This *in silico* study aims to motivate functional investigations that will consolidate the importance of this fascinating group of still poorly understood proteins in plants.

## Acknowledgements

We thank the International Research Training Group for Accelerating Crop Genetic Gain funded by The University of Queensland and the German Research Foundation (DFG) for enabling this study.

## CRediT authorship contribution statement

**Ian Godwin, Karen Massel and Gary Shenk:** conceptualization. **Carla Lopez:** writing – original draft, formal analysis, data curation. **Ian Godwin and Gary Shenk:** supervision, resources, project administration. **Marc Morris and Gary Shenk:** software, formal analysis, methodology, validation. **All authors**: writing – review and editing.

## Supporting information legends

**Figure S1.**
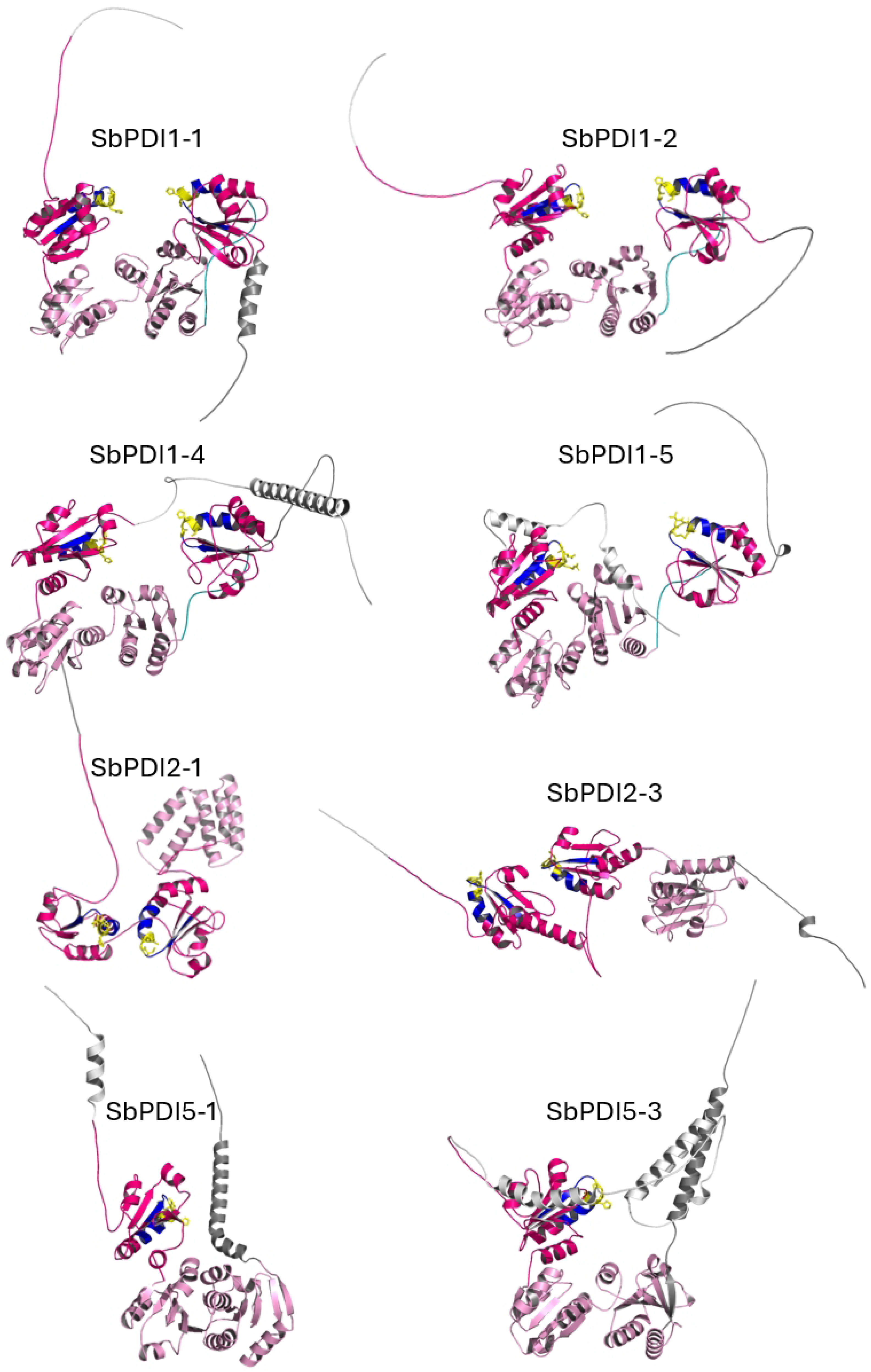
Predicted overall three-dimensional structures of SbPDIs. Note that SbPDI2-1b presents the same predicted structure as SbPDI2-1a. Detailed descriptions and comparisons follow in Results.

**Figure S2.**
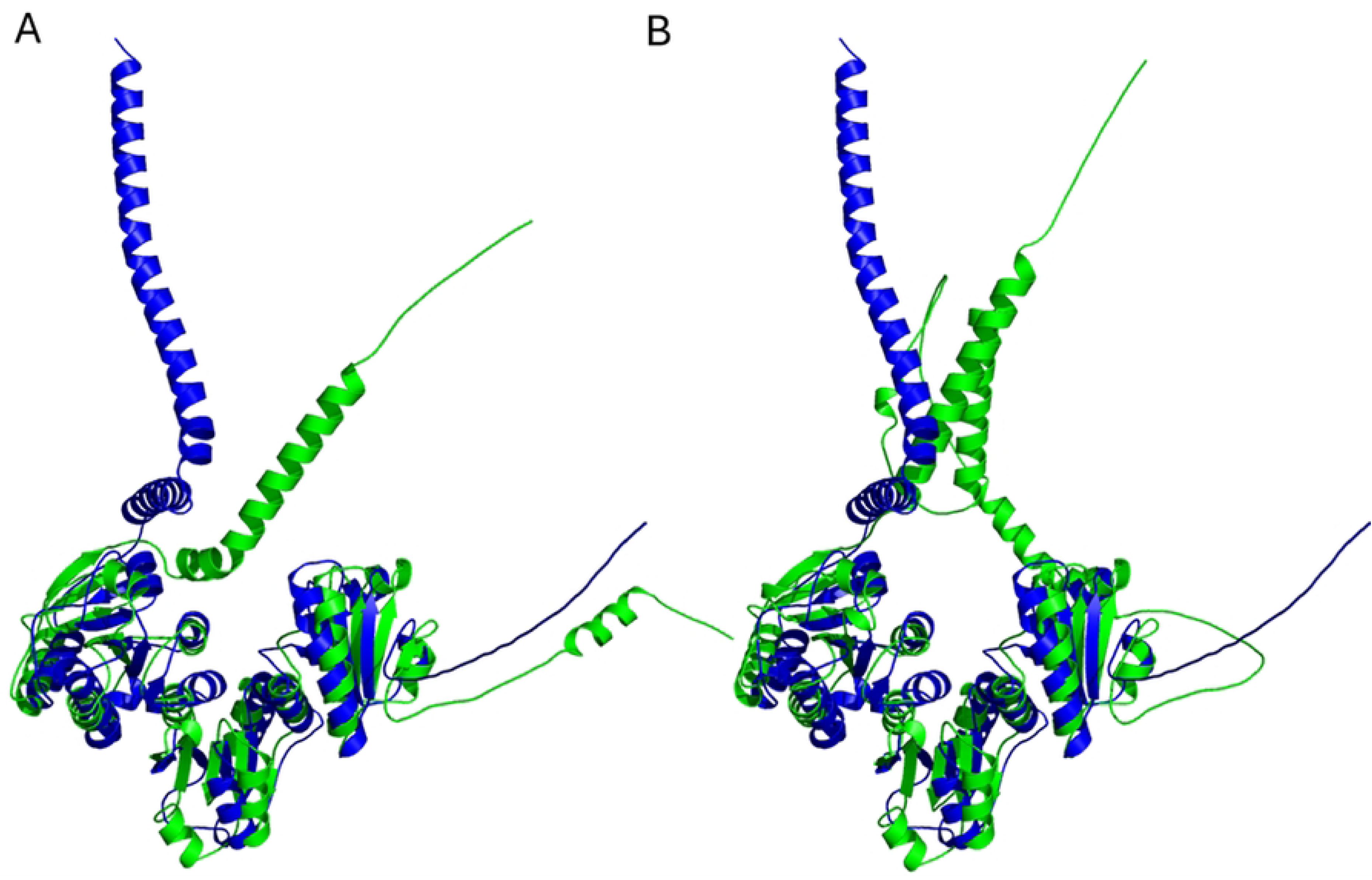
Superimposition of selected structural models of SbPDI5-2 and SbPDI5-3 with TMX3. (A) SbPDI5-2 (green) superimposes with an RMSD of 5.1 Å with the human TMX3 (blue). (B) SbPDI5-3 (green) greatly aligns with the TMX3 (blue) with an RMSD of 3.8 Å.

**Figure S3.**
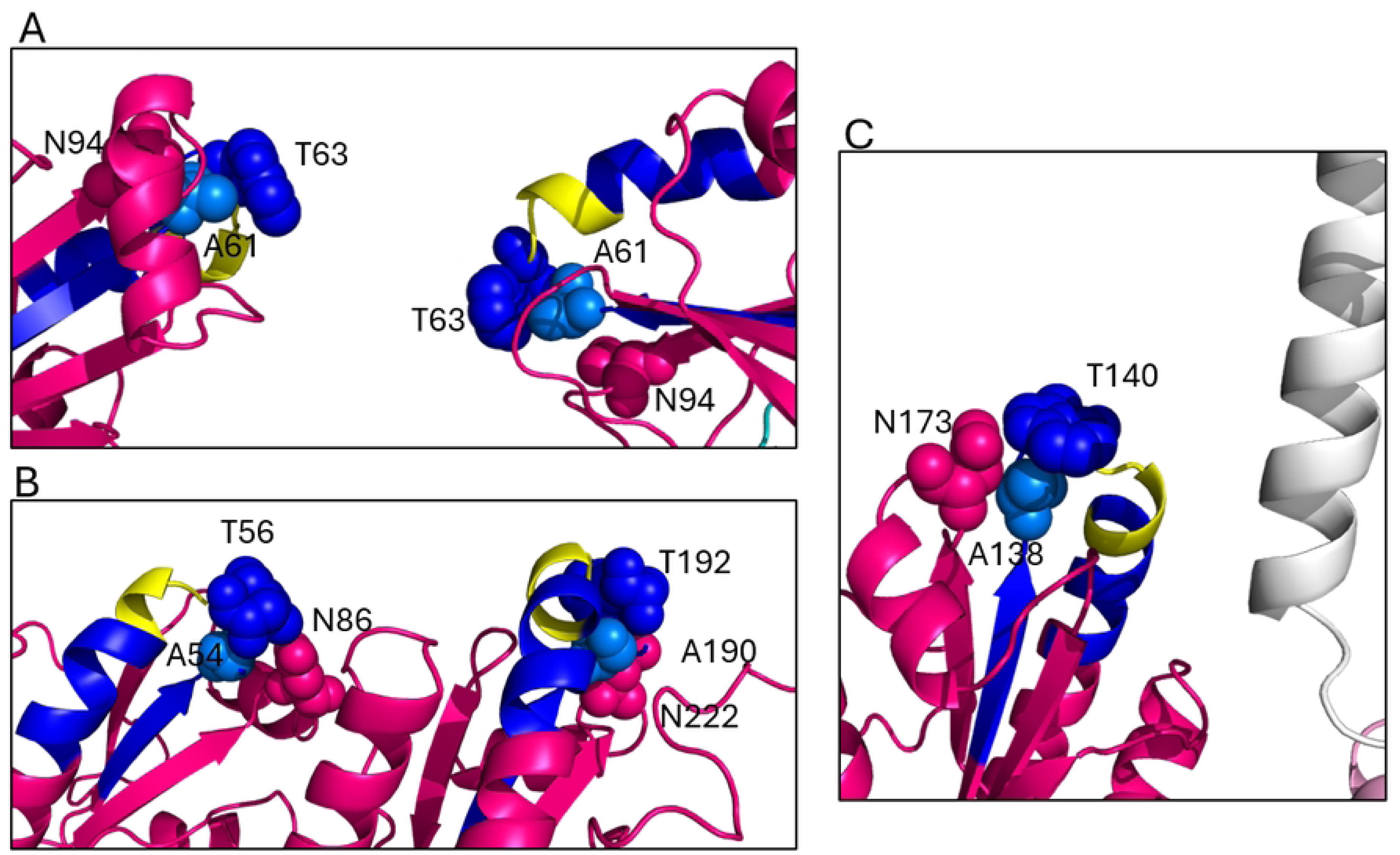
Position of residues that conform the structural cluster preceding the CXXC motif in SbPDI1-1 (A), SbPDI2-3 (B), and SbPDI5-3 (C). Residues are shown in spheres: tryptophan (blue), alanine (light blue), asparagine (pink).

